# ROOT PATTERNING AND REGENERATION ARE MEDIATED BY THE QUIESCENT CENTER AND INVOLVE BLUEJAY, JACKDAW AND SCARECROW REGULATION OF VASCULATURE FACTORS

**DOI:** 10.1101/803973

**Authors:** Alvaro Sanchez-Corrionero, Pablo Perez-Garcia, Javier Cabrera, Javier Silva-Navas, Juan Perianez-Rodriguez, Inmaculada Gude, Juan Carlos del Pozo, Miguel A. Moreno-Risueno

## Abstract

Stem cells are central to plant development. During root postembryonic development stem cells generating tissues are patterned in layers around a stem cell organizer or quiescent center (QC). How stem cell lineages are initially patterned is unclear. Here, we describe a role for BLUEJAY (BLJ), JACKDAW (JKD) and SCARECROW (SCR) transcription factors in patterning of cell lineages during growth and in patterning reestablishment during regeneration through regulation of number of QC cells and their regenerative capacities. In *blj jkd scr* mutants, QC cells are progressively lost which results in loss of tissue layers. Upon laser ablation *blj jkd scr* is impaired in QC division and specification resulting in severe impairment in pattern regeneration. Beyond direct regulation of QC activity by these transcription factors, reduced levels of SHORT-ROOT (SHR) and of PIN auxin transporters were observed in the vasculature of *blj jkd scr*, leading to strong reduction in the auxin response in the QC. We narrowed down non-cell-autonomous regulation of vascularly expressed genes in *blj jkd scr* to C-REPEAT BINDING FACTOR 3 (CBF3). *cbf3* mutant shows reduced levels of SHR in the vasculature, and in addition QC disorganization and downregulation of the QC regulator *WUSCHEL-RELATED HOMEODOMAIN 5 (WOX5)*. *CBF3* gene is primarily expressed in the ground tissue downstream of BLJ, JKD and SCR, while CBF3 protein may move. Targeted-expression of *CBF3* to the ground tissue of *blj jkd scr* recovers radial patterning and regeneration. We propose that BLJ, JKD and SCR regulate QC-mediated patterning, and that part of this regulation involves CBF3.

## INTRODUCTION

Plant developmental plasticity relies on meristems. During postembryonic development, meristematic cells are generated from the activity of stem cells [1]. Root stem cells have pre-assigned identities and thus they are understood as initial cells of the distinctive cell lineages of the root [2]. Stem cells and their lineages are organized in layers but the cues and developmental processes which pattern this radial organization are unknown. Stem cells are found in specific microenvironments called stem cell niches [3, 4]. Stem cell niches also contain organizer cells which promote stem cell activity and maintain surrounding cells in an undifferentiating state [5]. Organizer cells of the root stem cell niche are known as the quiescent center (QC) because of their slow division rate. QC divisions have been shown to be able to generate root stem cells during postembryonic development, and thus, the QC has been proposed to be a long-term reservoir for stem cells [6]. However, if formation of stem cells by the QC plays an active role during growth to generate a radial pattern, or it is rather an emergency mechanism in case a stem cell is accidentally lost is unknown.

Root stem cell niche specification and positioning in *Arabidopsis thaliana* requires hormone gradients and other signals, primarily converging into the PLETHORA (PLT) and SHORT-ROOT (SHR)/SCARECROW (SCR) pathways [3, 7-9]. Auxin hormone forms a gradient in the root meristem with a maximum in the stem cell niche [10]. PLT accumulation is interpreted as a slow-read of auxin maxima, and thus, a maximum in the accumulation of PLT proteins occurs in the stem niche area. PLT transcription factors (TFs) are mobile proteins, moving longitudinally in oppose direction to the auxin gradient. PLT function is dose dependent and high PLT concentration associates with stem cell niche specification and QC maintenance, whereas decreasing PLT concentrations may instruct cells to proliferate and finally differentiate [8, 11, 12]. SHR is also a mobile TF. *SHR* is transcriptionally expressed in the central vasculature cylinder and surrounding pericycle layer [13], although the mechanism controlling its transcription is not resolved. SHR protein moves radially to adjacent endodermis and QC, where it activates its targets SCR and the TF of the BIRD family JACKDAW (JKD) [14, 15] functioning as a dose dependent signal [16]. SCR and JKD interact with SHR and sequester it in nuclei. SHR regulates radial patterning being necessary to establish endodermis/cortex and metaxylem/protoxylem cell types [17-19]. SHR has also been reported to maintain and promote QC specific expression and activity in combination with SCR and JKD [3, 20]. SCR itself has been shown to regulate gibberellin and cytokinin signaling affecting auxin biosynthesis and root stem cell activity and, in addition it regulates cytokinin signaling from the endodermis which controls meristem size and root growth [21-23].

WUSCHEL-RELATED HOMEODOMAIN 5 (WOX5) is involved in specification of QC identity and in maintaining a low degree of differentiation of surrounding stem cells [5]. PLT, SHR, JKD and SCR TFs regulate *WOX5* specific expression in the QC [6, 8, 9, 20]. In addition, PLT and SCR interact with the TFs TEOSINTE-BRANCHED CYCLOIDEA PCNA (TCP) 20 and 21 to regulate *WOX5* expression through PLT binding sites in *WOX5* promoter [24].

QC divisions to replenish dead or wounded cells are critical for regeneration [25, 26]. QC quiescence is restricted by the RETINOBLASTOMA-RELATED (RBR) protein / SCR circuit and WOX5, working respectively, as repressors of mitotic transitions and CYCLIN D3;3/1;1 activity [6, 27]. Regenerative QC divisions can be induced upon supplementation with genotoxic agents that selectively kill stem cells or upon laser ablation of stem cells, showing that QC divisions are critical for stem cell niche regeneration [25, 26, 28]. These QC regenerative divisions are restricted by the RBR-SCR circuit, which is in turn is regulated by Jasmonic Acid. In addition, Jasmonic Acid activates ETHYLENE RESPONSE FACTOR (ERF) 109, ERF115 and CYCLIN D6;1 to promote regeneration [26, 28]. Removal of QC cells through laser ablation triggers the specification of a new stem cell niche involving WOX5, SCR and PLT as regulators of cell fate and auxin gradient formation [29].

Here, through analyses of *Arabidopsis thaliana* mutants in the transcription factors BLUEJAY (BLJ), JACKDAW (JKD) and SCARECROW (SCR) and chemical treatments, we show that QC cell number and their regenerative capacities maintain cell lineages and patterning downstream of these TFs. In addition, we show that these TF regulate specification of new QC-like cells upon laser ablation which is required to re-establish patterning during regeneration. Intriguingly, BLJ, JKD and SCR regulate PIN auxin transporters and SHR protein levels in the vasculature, although these TFs are expressed in the ground tissue and the QC. Decreased levels of SHR and auxin response were observed concomitant with loss of QC cells but prior major defects in radial patterning occurred. These results suggest the existence of non-cell autonomous regulation of vascularly expressed genes by BLJ, JKD and SCR, beyond direct regulation of QC gene expression. BLJ, JKD and SCR regulate the TF C-REPEAT BINDING FACTOR 3 (CBF3). CBF3, which was originally described as a cold response factor, shows specific expression in the ground tissue under standard temperature, while CBF3 protein is mobile. In the absence of BLJ, JKD and SCR, we detect loss of QC cells and decreased QC regenerative capacities, whereas targeted expression of *CBF3* to the ground tissue of *blj jkd scr* shows recovery of the radial patterning and stem cell niche regeneration. Therefore, CBF3 may contribute as a radial signal from the ground tissue to regulate patterning during development and regeneration.

## RESULTS

### BLUEJAY, JACKDAW and SCARECROW work combinedly to maintain root patterning and meristematic activity

Six members of the C_2_H_2_ TFs collectively known as the BIRDs have been shown to be involved in formation of the ground tissue lineage [15] as well as in delimiting boundaries between adjacent cell layers, such as the epidermis, endodermis and pericycle layers [18, 30]. Beyond known regulation of the ground tissue by this family of TFs, we hypothesized that the BIRDs might be involved in regulation of root patterning through specification and/or maintenance of tissue lineages and their stem cell initials.

There are thirteen members of the BIRD family of TFs. We decided to focus on JACKDAW (JKD) and BLUEJAY (BLJ) as downregulation of JKD through RNA interference showed disorganization of the stem cell niche [30] and we had observed reduced root growth in plants carrying the mutation *jackdaw-4 (jkd)* in combination with the mutation *bluejay (blj)* (Figure 1A). SCARECROW (SCR) is a GRAS TF involved in maintenance of stem cell and meristematic function and *scarecrow-4 (scr)* mutant has been shown to have reduced root growth [7]. Combination of *blj*, *jkd* and *scr* mutations showed additive defects, which resulted in almost root growth cessation from 8 days post imbibition (dpi) onwards (Figure 1A, S1A). Interestingly, our data show that *blj jkd scr* mutants continuously decreased growth while mutants in other regulators of stem cell and meristematic function such as *plethora (plt) 1-4* (*plt1*) *plt2-2* (*plt2*) mutant [9, 13] tended to maintain growth over all days analyzed; despite the fact growth is severely reduced in *plt1plt2*. Meristem inspection showed that although meristems of *blj jkd scr* mutants were of similar size as in the wild type (WT) in mature embryos (Figure S1B), by 6 dpi meristems of *blj jkd scr* were almost of similar size as *plt1 plt2* meristems, further reducing their size from 7 to 13 dpi (Figure S1C,D).

**Figure 1.**
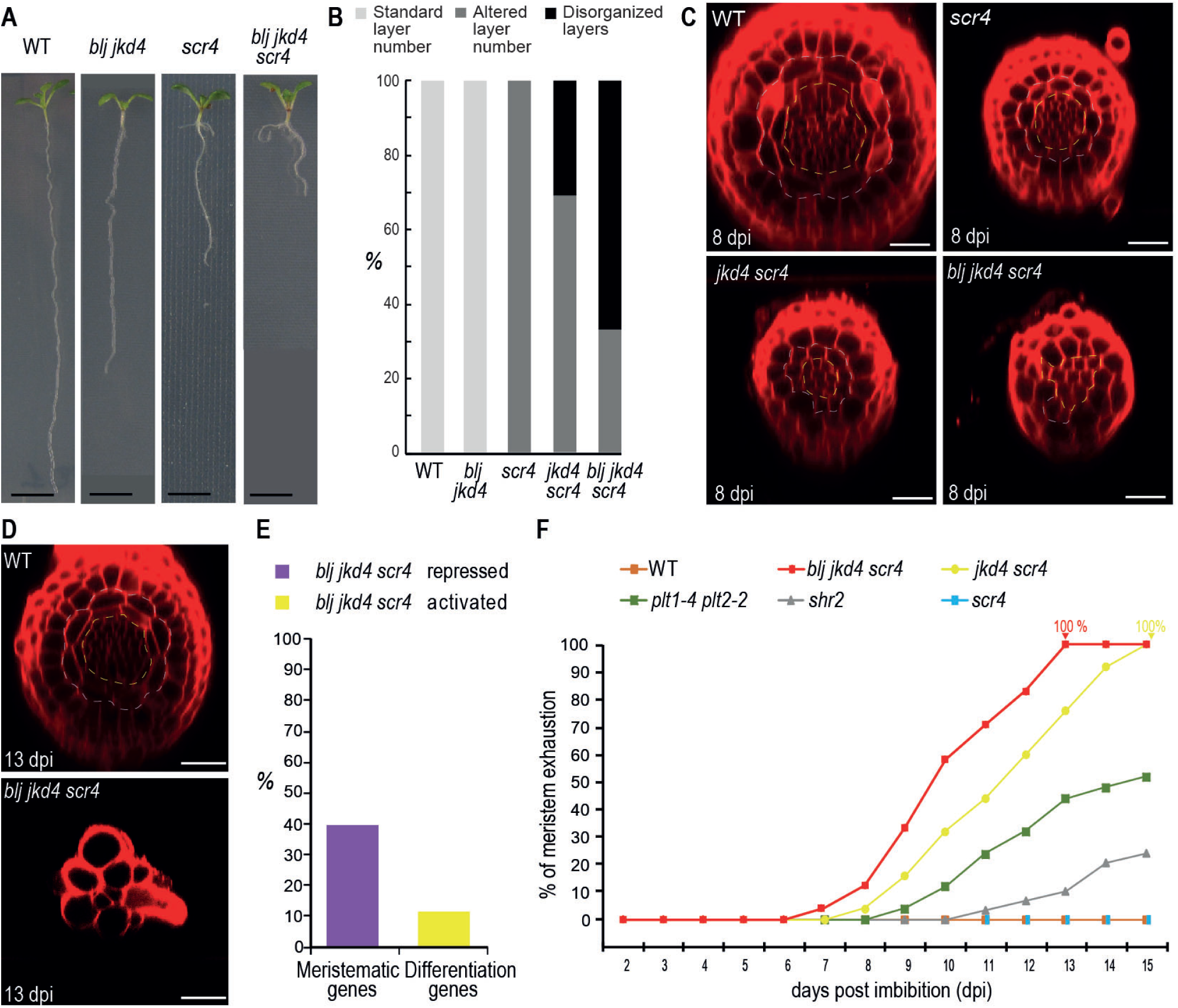
BLUEJAY, JACKDAW and SCARECROW work combinedly to maintain root patterning in Arabidopsis. **A**, images of seedlings of the wild type (WT) and of combinations of *jackdaw-4* (*jkd4)*, *scarecrow-4 (scr4)* and *bluejay (blj)* mutants at 10 days post imbibition (dpi). **B**, quantification of pattern alteration in WT, *blj jkd4*, *scr4*, *jkd4 scr4* and *blj jkd4 scr4* mutants at 8 dpi. Yellow dotted line: internal tissues (stele) boundary, blue dotted line: boundary between the ground tissue and the epidermis. **C**, confocal images showing representative radial patterns of WT, *scr4*, *jkd4 scr4* and *blj jkd4 scr4* plants at 8 dpi. **D**, confocal transverse sections of WT and *blj jkd4 scr4* meristems at 13 dpi**. A-D**, scale bar: 0.25 µm. **E**, percentage of differentiation-related or meristematic-related genes repressed or activated in microarrays profiling *blj jkd4 scr4* meristems prior major alterations in patterning were observed (5dpi). **F**, percentage of meristems showing root hairs (meristem exhaustion) in WT, *shortroot-2 (shr2)*, *scr4*, *jkd4 scr4*, *plethora1-4 (plt1-4) plt2-2*, and *blj jkd4 scr4*.

Analysis of *blj jkd scr* patterning showed profound alterations. Transverse meristem sections of *blj jkd scr* at 8 dpi showed no easily recognizable organization in layers with central cells showing palisade arrangement and reduced number of cells as compared to the WT (Figure 1B-C). In contrast, *scr* and most *jkd scr* mutants showed radial pattern organization in concentric layers at 8 dpi, although *jkd scr* also showed reduced number of cells in the root cylinder as *blj jkd scr*. In agreement with previous observations [15, 21], one ground tissue layers was observed for *scr* and *jkd scr* and none for *blj jkd scr* as compared with two layers observed in the WT. BLJ, JKD and SCR proteins accumulate in the ground tissue, and in addition, JKD and SCR proteins are present in the QC [15, 18]. As *blj jkd scr* mutants show disruption of the standard radial pattern while organization in concentric layers was still present in most *jkd scr* mutants, a non-cell autonomous mechanism regulating layer patterning from the ground tissue can be inferred. Furthermore, at 13 dpi only few enlarged cells made up the root cylinder of *blj jkd scr* (Figure 1D), indicating severe impairment of *blj jkd scr* meristems to pattern themselves and maintain cell lineages during postembryonic development. Because defects appeared from 6 dpi onwards, we report results in this paper between 6 and 13 dpi, normally at the first day it was detected.

### BLUEJAY, JACKDAW and SCARECROW regulate gene expression associated to proliferation and differentiation programs

To better understand defects in *blj jkd scr* we analyzed microarray experiments which profiled the *blj jkd scr* apical meristem at 5dpi [15], prior severe growth and patterning alterations were observed. Analysis of differentially expressed genes between wild type (WT) and *blj jkd scr* meristems showed repression of genes involved in regulation of cell cycle progression (Figure S2A) among others. Introgression of the G2 to M marker CYCB1;1 in *blj jkd scr* showed severe impairment in mitotic transition (Figure S2B), likely indicative of less cell proliferation rate. Further analyses of differentially expressed genes in the *blj jkd scr* apical meristem showed repression of about 40% of genes normally expressed in the meristem while over 10 % of genes normally expressed in the differentiation zone were activated in the meristematic region of this mutant (Figure 1E, Figure S2C-D). In agreement with this observation, we observed premature meristem exhaustion for *blj jkd scr* mutants between 8-13 dpi as compared to the WT, while it occurred later for *jkd scr* and did not occurred for *scr* (Figure 1F). Analysis of mutants in other factors involved in stem cell niche maintenance such as *short-root (shr)* and *plt1 plt2* showed reduced meristem exhaustion in comparison with *blj jkd scr* (Figure 1F), indicating that BLJ, JKD and SCR combined activity is critical for stem cell function. As stem cell activity and meristem differentiation is prevented by the QC, we hypothesized that QC function might be severely altered in *blj jkd scr*.

### BLUEJAY, JACKDAW and SCARECROW maintain number of quiescent center cells required for root patterning

To further investigate patterning defects observed in *blj jkd scr*, we introgressed QC and stem cell markers into this mutant. Our results show localization of the stem cell differentiation protein SOMBRERO (SMB) in cells located in the position of QC cells in *blj jkd scr*, whereas SMB is typically excluded from QC cells as observed in the WT (Figure 2A). Several cells in the position of the QC which appeared to have recently divided did not show localization of SMB (Figure 2A: inset), suggesting that more differentiated QC cells in *blj jkd scr* cannot undergo formative divisions. In agreement with SMB relocation, we observed that expression of the QC regulator *WUSCHEL-RELATED HOMEODOMAIN 5* (*WOX5*), which primarily locates in QC cells, appeared in reduced number of cells in *blj jkd scr* at 6 dpi as compared to the WT (Figure 2B). These results indicate reduced maintenance capacity of the QC in *blj jkd scr*, as shown by reduced number of QC cells which appeared to differentiate without undergoing formative divisions.

**Figure 2.**
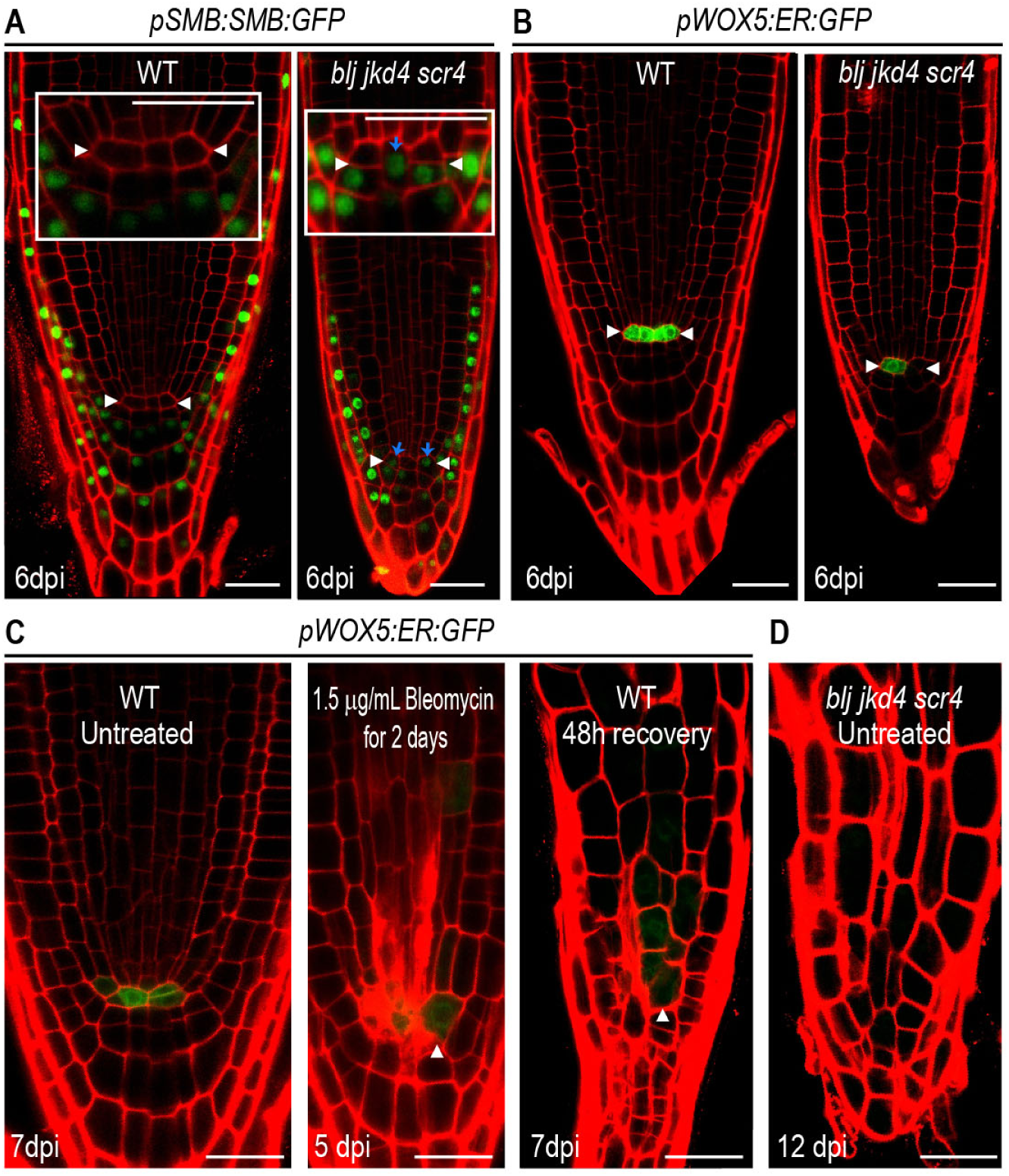
BLUEJAY, JACKDAW, SCARECROW maintain number of quiescent center cells required for root patterning. **A**, confocal images showing localization of SOMBRERO (SMB; *pSMB:SMB:GFP)*, in WT and *blj jkd4 scr4* meristems at 6dpi. SMB localizes in differentiating distal stem cells. Insets: close-up view of the stem cell niche in WT and *blj jkd4 scr4*. White arrowheads delimit the QC. Blue arrows: SMB localization in cells in position of the QC. **B**, Confocal images showing expression of *WUSCHEL-RELATED HOMEODOMAIN 5* (*WOX5*; *pWOX5:ER:GFP)* in WT and *blj jkd4 scr4* meristems at 6dpi. *WOX5* primarily marks quiescent center cells. White arrowheads delimit the QC. **C**, Confocal images showing expression of *WOX5* in WT control at 7dpi, in WT at 5 dpi after treatment with bleomycin and in WT after 48 hours of recovery after bleomycin treatment. White arrowhead: quiescent center cells and its lineage upon division. **D**, Confocal image of *blj jkd4 scr4* meristem at 12 dpi. **A-D**, Scale bars: 25 µm.

QC divisions have been shown to replace stem cells [6]. Reduced number of QC cells could result in less stem cell activity but also in less capacity of the QC to replace non-functional or damaged stem cells during development. To test this hypothesis, we treated WT roots with the genotoxic drug bleomycin. Bleomycin was shown to cause stem cell death [25, 26]. However, we observed that at higher concentration than reported, QC cells could be also damaged or killed. After two days of treatment, most QC cells were damaged in WT roots as indicated by propidium iodide staining inside the cell and absence of *WOX5* expression. The whole meristem appeared more differentiated showing enlarged cells, as expected from having reduced QC activity, but in addition, reduced number of vasculature layers was observed. Upon removal from treatment, remaining WT QC cells divided but failed to reconstitute a normal stem cell niche and meristem. Furthermore, these roots showed impaired pattering with reduced number layers (Figure 2C) which resembled *blj jkd scr* roots at 12 dpi (Figure 2D). This result indicates that reduced number of QC cells with reduced activity may result in altered patterning and be the cause of the observed defects in *blj jkd scr* mutant.

### BLUEJAY, JACKDAW and SCARECROW are required to maintain SHORT-ROOT and auxin transporter PIN-FORMED 1 levels in vascular tissues

QC specification requires the SHR/SCR pathway, particularly this regulation is carried out by SHR, SCR and JKD [24]. In addition to a direct effect on QC expressed genes caused by absence of JKD and SCR in *blj jkd scr*, our transcriptomic analysis showed that SHR regulations was also affected, as *SHR* gene was repressed in *blj jkd scr* (Figure S3A). To investigate SHR levels in *blj jkd scr*, we performed signal quantification using hybrid detector counting in confocal analyses as we had previously shown to accurately detect changes in gene expression and protein levels [31]. We observed less SHR accumulation in vascular tissues in *blj jkd scr* in comparison with the WT at 6 dpi (Figure 3A) whereas no obvious changes were detected at earlier days. Our results showed that reduction in number of QC cells and patterning defects in *blj jkd scr* aggravated from 6dpi onwards, which interestingly correlates with a five-fold decrease in SHR levels observed at 6 dpi.

**Figure 3.**
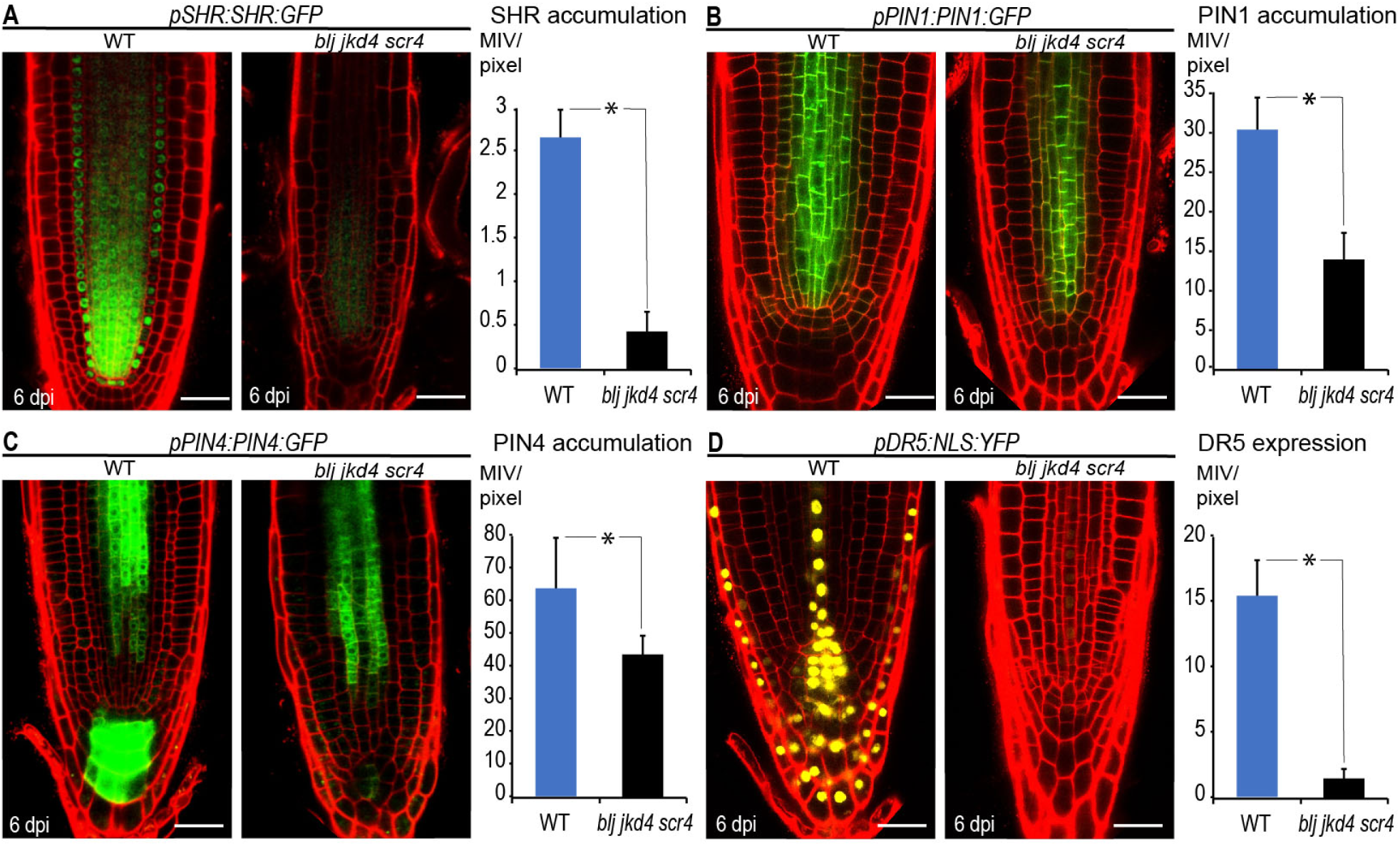
BLUEJAY, JACKDAW and SCARECROW are required to maintain SHORT-ROOT and auxin transporter PIN-FORMED 1 levels in vascular tissues. **A**, Confocal images showing accumulation of SHORT-ROOT (SHR, *pSHR:SHR:GFP*) in WT and *blj jkd4 scrc4* plants at 6dpi. Graph comparing SHR levels in WT and *blj jkd4 scrc4* plants at 6dpi. **B**, Confocal images showing accumulation of auxin transporter PIN-FORMED 1 (PIN1, *pPIN1:PIN1:GFP*) in WT and *blj jkd4 scrc4* plants at 6dpi. Graph comparing PIN1 levels in WT and *blj jkd4 scrc4* plants at 6dpi. **C**, Confocal images showing accumulation of PIN4 (*pPIN4:PIN4:GFP*) in WT and *blj jkd4 scrc4* plants at 6dpi. Graph comparing PIN4 levels in WT and *blj jkd4 scrc4* plants at 6dpi. **D**, Confocal images showing expression of the auxin response marker *pDR5:NLS:YFP* in WT and *blj jkd4 scrc4* plants at 6dpi. Graph comparing *pDR5:NLS:YFP* expression in WT and *blj jkd4 scrc4* plants at 6dpi. **A-D**, Scale bars; 25 µm. Asterisk: p-value<0.001 in a one-way ANOVA. MIV/pixel: mean intensity value/pixel.

QC specification has been also described as a slow readout of auxin maxima accumulation [11]. In order to investigate whether impairment in auxin gradient and maxima formation in *blj jkd scr* could also relate to the observed defects in the QC, we analyzed auxin efflux carrier expression. Our transcriptomic analyses showed a decrease in *PIN1*, *PIN2* and *PIN4* gene expression (Figure S3B). PIN1 and PIN4 are essential for formation of a functional auxin gradient with a maximum in the QC area [32]. Analysis of PIN1 and PIN4 accumulation in *blj jkd scr* showed decrease in the amount of PIN1 and PIN4 (Figure 3B, C). PIN1 levels decreased in vascular tissues while for PIN4 we primarily observed a strong reduction in the columella area. In agreement with a reduction in auxin transporters, we detected less auxin response in the QC of *blj jkd scr* as reported by the *DR5* marker (Figure 3D).

*SHR, PIN1* and *PIN4* genes are expressed in the vasculature and in addition, *PIN4* is expressed in the columella. In contrast, BLJ TF is specifically localized in the ground tissue, while JKD and SCR TFs accumulate both in the ground tissue and QC cells [15]. Based on reduced SHR and PIN levels outside of the ground tissue and QC cells in *blj jkd scr*, a non-cell-autonomous effect is observed in the regulation carried out by BLJ, JKD and SCR.

### Identification of ground tissue transcription factors which might be mobile

To investigate non-cell-autonomous regulation in BLJ, JKD and SCR activities we focused on TFs, as mobile TFs have been shown to mediate non-cell-autonomous regulation during development. In addition, BLJ, JKD and SCR regulate most SHR responses in the ground tissue [15], so for searching purposes we filtered TFs in the SHR network, with enriched expression in the ground tissue (Figure 4A). Next, we tested if the identified TFs may move to adjacent tissues. The J0571 enhancer is expressed in the ground tissue and occasionally in QC cells, and it can be used to drive gene expression to these tissues using the UAS promoter. Thus, we used the J0571*/GAL4-UAS::TF* system to direct expression of the identified TFs. These TFs were also tagged with a yellow fluorescent protein (YFP). Out of the TF tested (Supplementary Table 1), three TFs: C-REPEAT BINDING FACTOR 3 (CBF3), DUO1-ACTIVATED ZINC FINGER 3 (DAZ3) and NUCLEAR FACTOR Y SUBUNIT B5 (NF-YB5) showed YFP signal outside of the ground tissue/QC transcriptional domain (GFP marked), indicating transcript or protein movement (Figure 4B, Figure S4A, B). Rest of TF tested using the same system did not show YFP signal outside of the ground tissue or the QC (Supplementary Table 1, Figure S4C,D) indicating that gene expression levels driven by this system is not promoting *per se* protein or transcript movement. Interestingly, a radial pattern with extra divisions in the ground tissue was observed for J0571*/GAL4-UAS::*CBF3, J0571*/GAL4-UAS::*DAZ3 and J0571*/GAL4-UAS::*NF-YB5.

**Figure 4.**
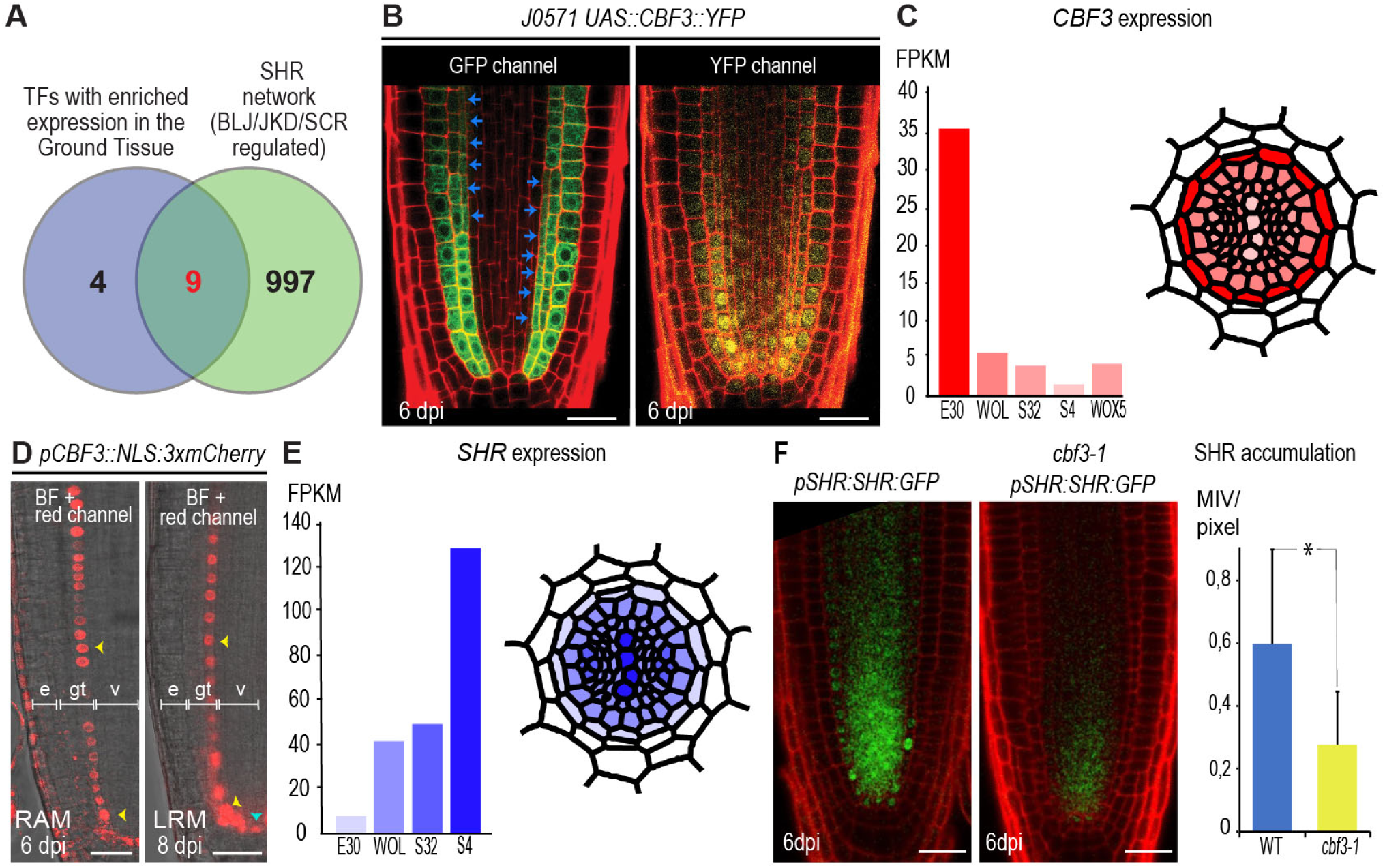
C-REPEAT BINDING FACTOR 3 is a transcription factor downstream of BLUEJAY, JACKDAW and SCARECROW which regulates SHORT-ROOT levels in the vasculature. **A**, venn diagram. **B**, confocal images showing localization of C-REPEAT BINDING FACTOR 3 (CBF3) tagged with a yellow fluorescet protein (YFP) in the UAS/GAL4 system using the enhancer trap line J0571, which primarily drives expression to the ground tissue (GFP marked) and ocassionaly to several QC cells. Blue arrows: extra divisions in the endodermis. **C**, transcript levels of *CBF3* as reported by RNA sequencing in the high-resolution expression map of the Arabidopsis root [33]. E30: endodermis, WOL: stele, S32: protophloem, S4: xylem, WOX5: QC. The cartoon represents a root transversal section colored with *CBF3* expression as in the graph. **D**, *CBF3* promoter expression in the root apical (RAM) and lateral root (LRM) meristems. Arrowheads: nuclear localized mCherry fluorescent protein under the control of *CBF3* promoter in the endodermis (yellow arrowheads) or the QC (blue arrowheads). e: epidermis, gt: ground tissue, v:vasculature. **E**, transcript levels of *SHR* as reported by the high-resolution expression map of the Arabidopsis root. The cartoon represents a root transversal section colored with *SHR* expression as in the graph. **F**, confocal images of *pSHR:SHR-GFP* (SHR) in the RAM of WT and *cbf3-1* at 6 dpi. Graph comparing the accumulation of SHR in the RAM of WT and *cbf3-1* at 6 dpi. Asterisk: p-value<0.01 in a one-way ANOVA. MIV/pixel: mean intensity value/pixel. **B,D,F**, Scale bars: 25 µm.

### *C-REPEAT BINDING FACTOR 3* is a transcription factor with enriched expression in the endodermis which regulates SHORT-ROOT levels in the vasculature

Out of the three putatively mobile TF we identified, we focused on CBF3 because protein levels in vascular tissues for J0571*/GAL4-UAS::CBF3::YFP*, based on YFP signal, were more prevalent than for DAZ3 and NF-YB5 (Figure 4B, Figure S4A, B). In addition, we observed greater number of divisions in the ground tissue for J0571*/GAL4-UAS::CBF3::YFP* which occasionally created an extra layer. Based on a high-resolution expression map of the Arabidopsis root performed through RNA sequencing [33], *CBF3* is shown to have enriched expression in the endodermis while its expression levels in vascular tissues would be reduced by 7-fold (Figure 4C). To further explore *CBF3* expression pattern we fused the fluorescent mCherry protein to a nuclear localization signal (NLS) and express this reporter gene under the control of the 3 kb upstream promoter of *CBF3*. Confocal analyses showed enriched expression in endodermis for the nuclear localized mCherry under the *CBF3* promoter. In addition, expression driven by the 3 kb *CBF3* promoter could also be detected in the endodermis, several cortex cells and the QC of lateral roots (Figure 4D). Although *CBF3* expression associates with cold response [34], we found that *CBF3* was expressed in the endodermis under standard growth conditions (22 °C).

*SHR* gene has been reported to be expressed in vascular tissues [14], and accordingly the high-resolution expression map of the Arabidopsis root [33] showed enriched expression in the stele, primarily in xylem tissues (Figure 4E). To investigate if CBF3 may regulate SHR levels we quantified *pSHR::SHR::GFP* signal using hybrid detector counting in confocal analysis and observed reduced SHR protein accumulation in *cbf3* mutant (Figure 4F). As *CBF3* and SHR transcripts in vascular and ground tissue cell types show complementary expression patterns (Figure 4C-E) *SHR* regulation by CBF3 might involve CBF3 movement to the vasculature.

### C-REPEAT BINDING FACTOR 3 regulates quiescent center organization and root patterning downstream of BLUEJAY, JACKDAW and SCARECROW

Next, we investigated if *CBF3* was functionally regulated by BLJ, JKD and SCR at 6 dpi, as at that developmental time we had observed reduced number of QC cells in *blj jkd scr*. RT-PCR analysis of *CBF3* expression in *blj jkd scr* showed downregulation at 6 dpi (Figure 5A). To investigate if CBF3 may regulate stem cell niche organization, we analyzed *cbf3* loss of function mutant and *35S::CBF3* overexpressing line. We observed altered organization of the stem cell niche in *cbf3*, which showed QC cells with abnormal morphologies (penetrance 35 %). In contrast, *35S::CBF3* stem niche was always well organized and tended to show two QC cells at the stem cell niche middle plane while the WT normally has three at 9dpi (Figure 5B). To assess if CBF3 may regulate functionality of the stem cell niche, we studied expression of the QC regulator WOX5. We observed decreased expression of *WOX5* in *cbf3*, which correlated with QC disorganization (Figure 5C-D). Reduced *WOX5* expression can be indicative of a QC with reduced functionality resulting in a more differentiated stem cell niche. In contrast, *CBF3* overexpressing line showed increased *WOX5* expression in comparison with the WT.

**Figure 5.**
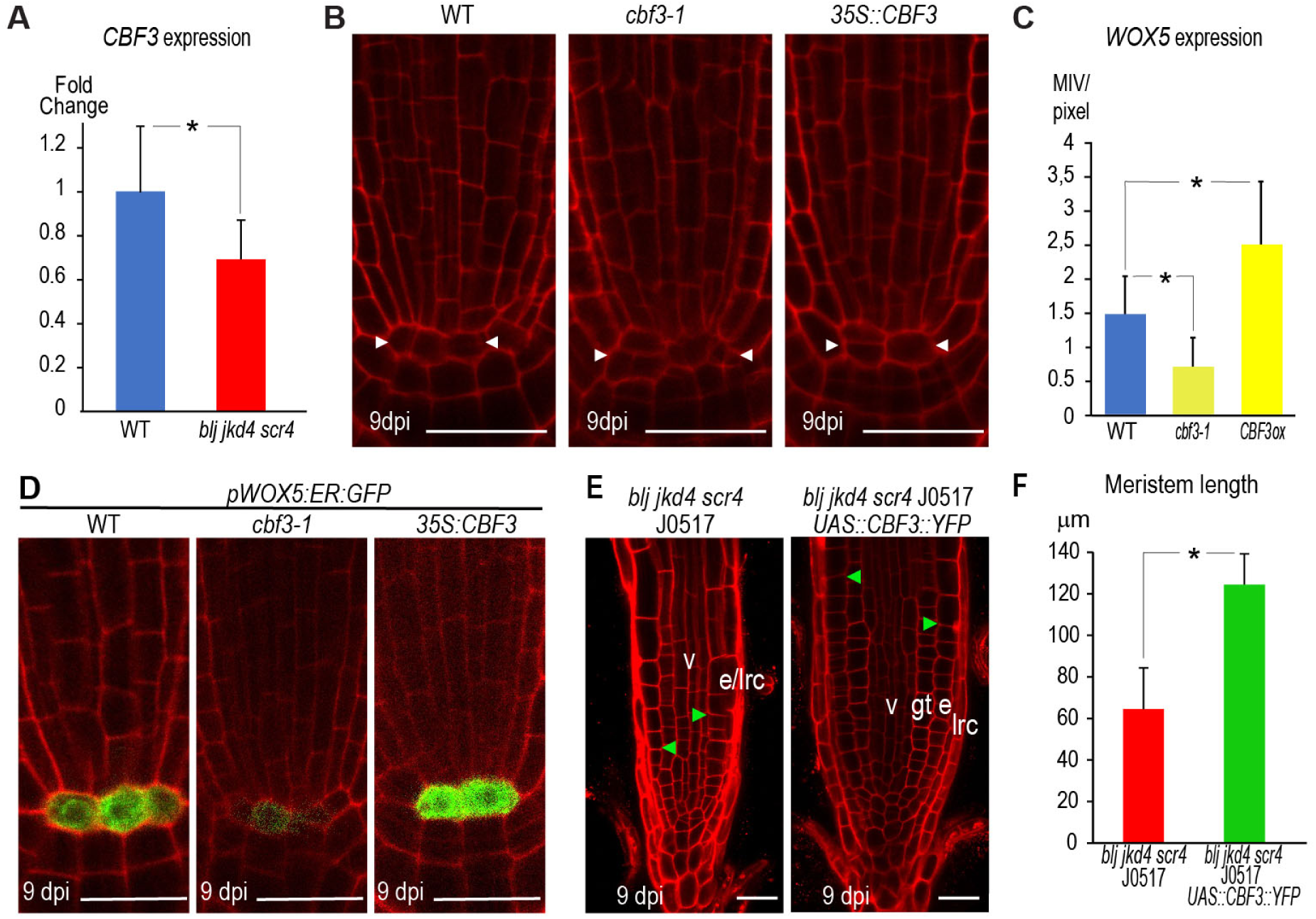
C-REPEAT BINDING FACTOR 3 regulates quiescent center organization and root patterning downstream of BLUEJAY, JACKDAW and SCARECROW. **A**, Real-time PCR analysis of *CBF3* expression in WT and *blj jkd4 scr4* RAMs at 6 dpi. Asterisk: p-value < 0.05 in a one-way ANOVA analysis. **B**, confocal images of the stem cell niche of WT, *cbf3-1* and p35S:CBF3 at 9dpi. White arrowheads delimit the QC. **C-D**, Quantification (**C**) of confocal images (**D**) of *pWOX5:ER:GFP* in WT, *cbf3-1* and p35S:CBF3 plants at 9dpi. Asterisk: p-value < 0.01 by General Linear Model (GLM) and LSD post-hoc test as compared to the WT. MIV/pixel: mean intensity value/pixel. **E**, confocal images of meristems of *blj jkd4 scr4* J0571 and *blj jkd4 scr4* J0571 introgressed with *UAS:CBF3:eYFP* at 9 dpi. lrc: lateral root cap, e: epidermis, gt: ground tissue, v:vasculature. Green arrowheads: end of the meristem. **F**, Quantification of meristem sizes of *blj jkd4 scr4* J0571 and *blj jkd4 scr4* J0571 *UAS:CBF3:eYFP* at 9dpi. Asterisk: p-value<0,001 in a one-way ANOVA. **B,D-F**, Scale bars: 25 µm.

As CBF3 was under the control of BLJ, JKD and SCR and was necessary for SHR and QC organization, we investigated if CBF3 could regulate root patterning from the BLJ, JKD and SCR transcriptional domain. We introgressed J0571*/GAL4*-*UAS::CBF3::YFP* into *blj jkd scr*. We observed increased meristem size as compared to *blj jkd scr* J0571, and notably, recovery of the radial pattern organization with primary tissue layers being recognizable, while no obvious radial pattern could be easily identified for *blj jkd scr* (Figure 5E-F). In contrast, when we introgressed the J0571*/GAL4-UAS::DAZ3* and *J0571/GAL4-UAS::NF-YB5* lines in *blj jkd scr* we observed modest or no recovery (Figure S4E), indicating that the observed effects were unlikely to be caused by expression driven by the J0571*/GAL4*-*UAS* system itself but that it was dependent on *CBF3*.

### Quiescent center formative divisions and stem cell niche regeneration requires BLUEJAY, JACKDAW and SCARECROW

Our results show that BLJ, JKD and SCR were required to maintain number of QC cells capable of undergoing formative divisions during development. To further investigate if QC cells in *blj jkd scr* could undergo formative divisions involved in stem cell regeneration, we treated roots with bleomycin using a concentration which specifically kills stem cells (0.8 μg/mL). After 24-hour treatment we observed reduced number of dead cells in *blj jkd scr*, suggesting reduced numbers of stem cells in this mutant as compared to the WT. Notably, upon bleomycin recovery, we detected few QC divisions in *blj jkd scr*, while QC cells in the WT actively divided as could be visualized by WOX5-GFP (Figure 6A).

**Figure 6.**
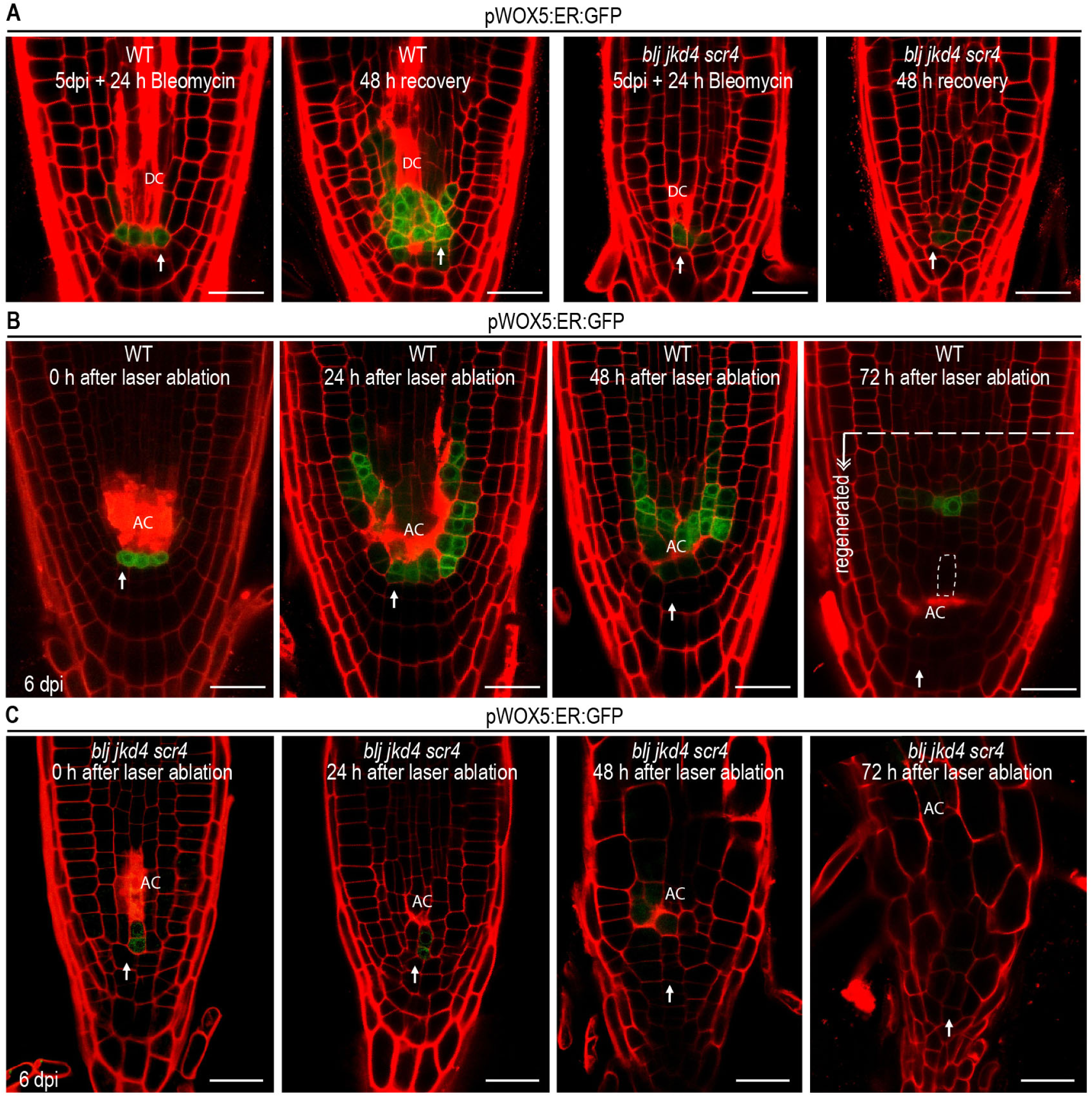
Quiescent center formative divisions and stem cell niche regeneration require BLUEJAY, JACKDAW and SCARECROW. **A**, confocal 48-hours (h)-time-lapse images of stem cell niche regeneration upon low dose of bleomycin treatment (0.8 ug/mL) killing stem cells. WT and *blj jkd4 scr4* roots carrying the *pWOX5:ER:GFP* QC marker were treated for 24h and then moved to bleomycin-free medium to asses QC divisions. **B-C**, confocal 72h-time-lapse images of stem cell niche regeneration upon laser ablation of stem cells above the QC, in WT and *blj jkd4 scr4* roots carrying the *pWOX5:ER:GFP* QC marker. AC: ablated cells. Dashed line: elongated cells (twice the size than adjacent upper cells) following regeneration. **A-C**, White arrow: a cell adjacent to the QC followed up during the regeneration process. Scale bars: 25 µm.

To test whole stem cell niche regeneration in *blj jkd scr*, we performed laser ablation of stem cells above the QC, as this procedure results in the re-specification of a new stem cell niche. Ablated cells collapsed over time forming a scar, while division of QC cells occurred concomitant with specification of newly WOX5-GFP marked cells for first 48 hours following ablation. In *blj jkd scr*, QC cells divided less than in the WT and in addition there was a strong reduction in the number of newly specified WOX5-GFP marked cells (Figure 6B-C, Figure S5A). At 72 hours after ablation, WOX5-GFP expression in the WT was confined to a few cells (the new QC) within the new stem cell niche, whereas cells more distal to the new QC, and adjacent to the scar, started to elongate resembling columella differentiation. In contrast, no regeneration of a new stem cell niche was observed in *blj jkd scr* meristems which degenerated (Figure 6C).

### Re-establishment of the radial pattern during regeneration involves *CBF3* expression in the ground tissue

To determine the role of CBF3 in stem cell niche regeneration, we performed laser ablation in J0571/GAL4-UAS::CBF3::YFP introgressed into *blj jkd scr* as well as in *cbf3* and *35S::CBF3*. Root regeneration rate in *blj jkd scr* mutant was severely compromised in agreement with our results (Figure 7A, 6B-C). Strikingly, we detected that regeneration capacity of *blj jkd scr* expressing J0571/GAL4-UAS::CBF3::YFP occurred at similar rates as the WT. When we analyzed radial pattern re-establishment during regeneration, we found that those roots of *blj jkd scr* J0571 which occasionally regenerated (~5%), were incorrectly patterned showing disorganized layers. In contrast, introgression of *CBF3* in *blj jkd scr* using J0571/GAL4-UAS::CBF3::YFP allowed pattern re-establishment, which was similar to the radial pattern observed for the WT (Figure 7B). As J0571 occasionally drives expression to several QC cells, it is possible that pattern regeneration requires certain levels of *CBF3* in the QC in addition to *CBF3* being expressed in the ground tissue.

**Figure 7.**
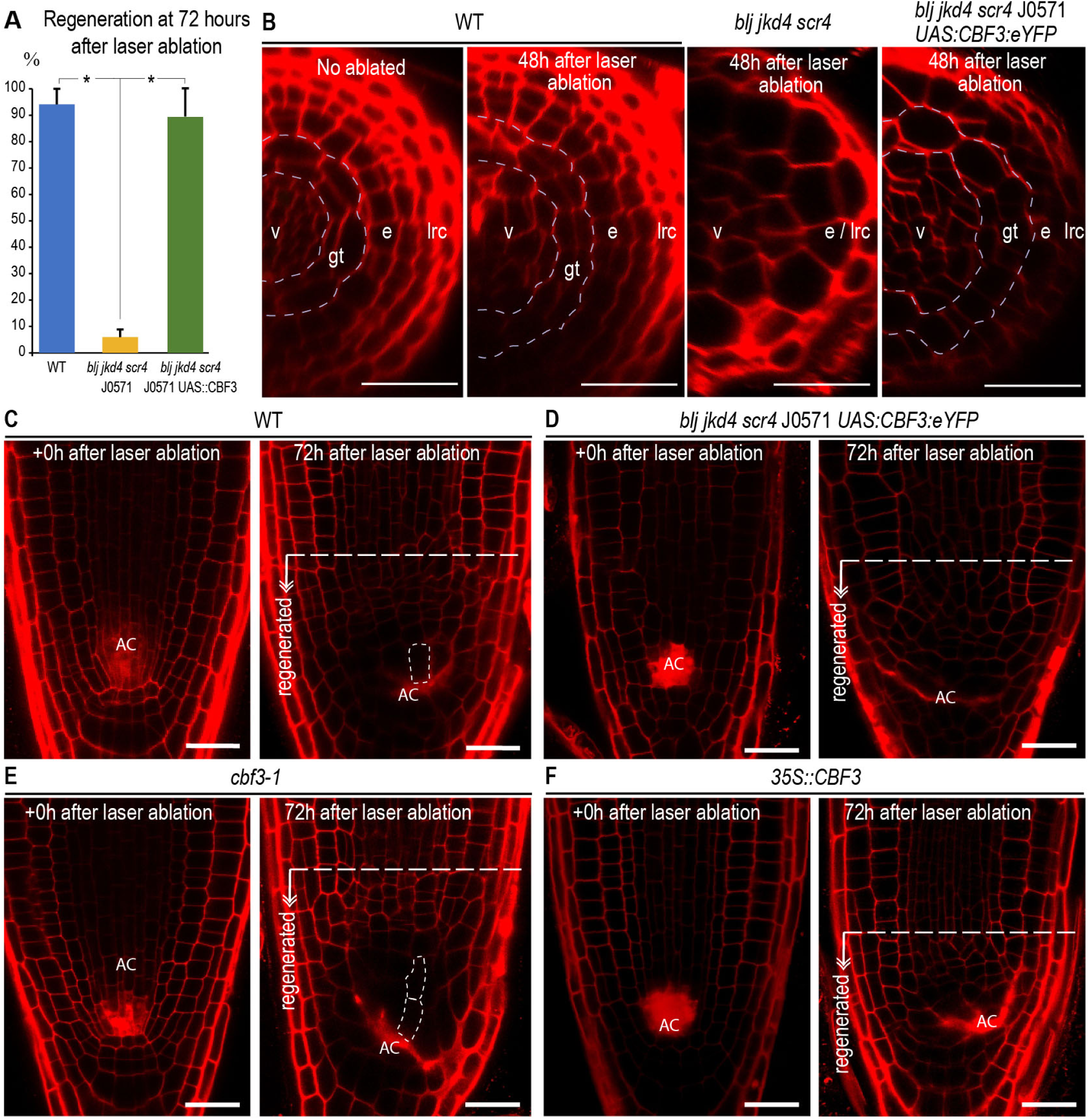
CBF3 regulates root patterning and stem cell niche specification during regeneration. **A**, graph showing percentage of regeneration following laser ablation in WT, *blj jkd4 scr4* J0571 and *blj jkd4 scr4* J0571 introgressed with *UAS::CBF3::YFP*. Asterisk: p-value < 0.001 by GLM and LSD post-hoc test. **B**, confocal images of transversal meristem section at 48h after laser ablation. lrc: lateral root cap, e: epidermis, gt: ground tissue, v: vasculature. Dashed blue line: contour delimiting the ground tissue. **C-F**, confocal images of stem cell niche regeneration upon laser ablation of cells above the QC for WT, *blj jkd4 scr4*, *blj jkd4 scr4* J0571 *pUAS:CBF3:eYFP*, *cbf3-1* and *35S::CBF3*. AC: ablated cells. Dashed line: elongated cells (twice the size than adjacent upper cells) following regeneration. **B-F**, Scale bars: 25 µm.

Evaluation of regenerated stem cell niches in *blj jkd scr* J0571/GAL4-UAS::CBF3::YFP revealed absence of elongated cells following regeneration, whereas these type of cells were observed in the WT (Figure 7C, D). Regenerated stem cell niches of *35S::CBF3* did not show elongated cells either, whereas *cbf3* regenerated stem cell niches tended to have two rows of elongated cells (Figure 7E, F). As cell elongation associates with differentiation, these results indicate that CBF3 also regulates the differentiation state of the stem cell niche during regeneration.

## DISCUSSION

Over last decades organ formation and differentiation have been shown to rely on positional information delivered by hormone gradients and mobile TFs. In the root meristem, SHR movement from inner vasculature to the ground tissue was shown to regulate important aspects of radial tissue organization in layers or radial patterning from preexisting cell lineages, such as cortex/endodermis specification from the ground tissue initial and protoxylem/metaxylem specification from vascular initials [13, 14]. How initial cell lineages are patterned is unknown. Our work provides evidences on how cell lineages are patterned from formative divisions of the stem cell organizer or QC during development. In addition, we show how these divisions require two homologous factors of the BIRD family, BLJ and JKD in combination with the GRAS TF SCR.

### Cell-autonomous and non-cell-autonomous regulation of quiescent center formative divisions mediating root patterning

The QC was originally shown to maintain surrounding stem cells active and undifferentiated [5]. More recently, the QC has been shown to be able to replace damaged stem cells upon stress [25, 26] and during development [6]. However, the functional implications of stem cell replacement by the QC, i.e. in root patterning, are missing. Our results indicate that reduced number of QC cells with low regenerative capacities (as in *blj jkd scr* mutant or wild-type bleomycin-treated roots) results in loss of cell lineages. Based on these results, the QC emerges as critical component in root patterning, constituting thus, as an active source of stem cells and/or cell lineage initials during development. As root stem cells are thought to have self-renewing properties and divisions of the QC to replace damage stem cells are not often observed, different scenarios can be envisioned. One possibility is that stem cell loss during development is not rare. As loss of the initial means missing the entire lineage, profound patterning alterations would be observed even if just one initial was lost every 1-2 days without replacement. Other possibility is that not all tissue initials had self-renewing capacity (or this is very low), relying likewise in QC formative divisions to replace them and maintain cell lineages. In this aspect, cell lineages firstly lost in *blj jkd scr*, such as the cortex/endodermis initial, could correspond to stem cells with low self-renewing capacity. In agreement with this possibility, the cortex/endodermis initial was observed to undergo formative periclinal divisions without apparently self-regenerating in 5-day-old roots [35].

JKD and SCR have been shown to cell autonomously regulate QC specification [20, 24]. In addition, our data reveal that these two TFs, in combination with BLJ which is not present in the QC, are required for QC formative divisions. Regulation of QC formative divisions might therefore be cell-autonomous regulated by JKD and SCR. In addition, non-cell autonomous regulation is required as highlights the fact that *blj jkd scr* shows loses organization in concentric layers whereas most *jkd scr* mutants still have a recognizable pattern. In our model, CBF3, primarily expressed in the ground tissue, emerges as a component in the regulation of vascularly expressed factors leading to QC formative divisions and stem cell specification such as SHR (as shown by less SHR in *cbf3)*. Furthermore, expressing *CBF3* (*J0571 UAS::CBF3::YFP*) in *blj jkd scr* ground tissue (and occasionally in the QC) rescues many of the patterning defects observed in this mutant, which confirms the importance of non-cell-autonomous regulation in cell lineage specification and organization.

### A plausible model for coordination of stem cell niche specification in the root meristem

In addition to JKD and SCR, SHR movement to the QC is also required for stem cell niche specification [14, 18]. In the longitudinal axis additional positional information to specify the QC and the stem cell niche is delivered by auxin hormone gradient. Auxin maxima induce PLT and high PLT dose associates with stem cell niche specification [11]. Based on our data, CBF3 is a new component in the regulation of positional information in the root meristem. Coordination between SHR and the auxin gradient to specify the stem cell niche has not been previously demonstrated. Our results show that BLJ, JKD and SCR and their downstream regulated factor CBF3, which are in the SHR pathway, regulate SHR levels (as shown by less SHR in *blj jkd scr* and *cbf3).* In addition, we show that BLJ, JKD and SCR regulate auxin transport and auxin gradient formation, which might be exerted through CBF3 as suggested by recovery in *blj jkd scr* meristem size when *CBF3* was expressed in the ground tissue of *blj jkd scr (blj jkd scr J0571 UAS::CBF3::YFP* line). In this scenario changes in the levels of SHR (or its downstream targets BLJ, JKD and SCR) could modify auxin levels through CBF3, in this was coordinating SHR and auxin pathways.

### Regulation of genes required for stem cell niche specification and regeneration involves CBF3

SHR and its direct target JKD were shown to have a specific interaction that promoted *WOX5* activation; and accumulation of JKD in the QC area was sufficient to induce specification of QC-like cells [20]. SCR was shown to be required for specification and maintenance of the QC [21], and more recently, it has been shown to work in combination with PLT3 and TCP20 to activate *WOX5* expression [24]. In this model of regulation, SHR and JKD on the one hand, and PLT and TCP, on the other, emerge as critical regulators of gene transcription leading to QC specification, while SCR might regulate their combinatorial interactions or be a scaffold. Our data show that CBF3 may promote SHR accumulation, which in agreement with these results, could lead to QC specification. Intriguingly, we detected recovery in stem cell niche specification and complementation in the absence of JKD and SCR in the QC (*blj jkd scr* mutant), suggesting that an increase in SHR levels might compensate for absence of these two factors. An alternative hypothesis might involve CBF3 directly binding to genes involved in QC specification and formative divisions to activate their transcription.

### Patterning re-establishment during other regenerative processes might involve CBF3

Pattern restoration upon cell-type targeted ablation or wounding has been shown to involve SHR and PLT triggering formative divisions in inner adjacent cells [36]. As SHR moves from inside to outside, signaling and activation of regenerative divisions from wounded cells appears to require the existence of unknown positional signaling. Our data show that *CBF3* expression in the ground tissue (which moves from outside to inside) is sufficient to restore stem cell regeneration upon ablation of specific vasculature cells. Future research might reveal a role for CBF3 as a signal delivering positional information required for cell fate specification upon wounding.

## MATERIALS AND METHODS

### Growth Conditions

Seeds were sterilized in 25% (m/v) NaClO and rinsed three times with sterile water before being sowed on 120 × 120 × 10 mm petri dishes containing 65 mL of one-half-Murashige & Skoog (MS) medium with 1% Sucrose and 10 g/L Plant Agar (Duchefa). For all the experiments, after 2 days of stratification at 4°C in darkness, plates were transferred to a custom made growth chamber and maintained vertically at 22°C with 16/8 hours light/darkness photoperiod at a white light rate intensity of 50 µmol·m^−2^·s^−1^. For bleomycin treatment, seeds were sown on MS medium with 0.8 μg/mL or 1.5 μg/ml bleomycin sulfate (Calbiochem) as indicated.

### Generation of constructs

For generation of pCBF3:NLS:3xmCherry line, a fragment of 2776 bp upstream of *CBF3* start codon was amplified by PCR (Phusion High-Fidelity DNA Polymerase Thermo Fisher Scientific) using F 5′-ACTTCTTTGCTTCACATAAGTTAAAAGTCA-3′and R 5′-TCTTGAAACAGAGTACTCTTCTGATCA-3′ primers from *A. thaliana* (Col-0) genomic DNA. The fragment was cloned into pNYZ28-A (NYZ tech) and P4P1R entry vector. For generation of UAS constructs, the UAS promoter [21] was cloned into P4P1R entry vector and coding sequences amplified by PCR with primers *CBF3-* At4g25480 F 5′-CACCATGAACTCATTTTCTGCTTTTTC-3′ and R 5’-ATAACTCCATAACGATACGTCG-3’, NF-YB5-AT2G47810 F 5’-CACCATGGCGGGGAATTATCATTCG-3’ and R 5’-ATTATCTGGCGAGGATTTAGG-3’, BHLH114/E29 (At4g05170) F 5’-CACCATGAACATGCATAATAGTTTCTTC-3’ and R 5’-CAAAGAATGGAAGATGCCAGA-3’, BHLH103/B70/E30 (At4g21340) F5’-CACCATGACAGAAGAATTCGACACTA-3’ and R 5’-ATTCCACAAACTAGATGTGTC-3’ and DAZ3 (At4g35700) 5’-CACCATGAGTAATCCCGAGAAGTCT-3’ and R 5’-CTCGACTTTATCATCGTTCTC-3’ and cDNA as template and introduced into pD-TOPO entry vector. Final constructs into dpGreenBarT vector were generated using 3-fragment GATEWAY System (Thermo Fisher Scientific) and transformed into Col-0 or J0571 as indicated using floral dipping method.

### Plant Material

Columbia-0 (Col-0) and Wassilewskija-2 (Ws-2) accessions were used as a genetic background as corresponding. The reporter lines pCYCB1;1:CYCB1;1:GFP [37], *shr-2* pSHR2:SHR2:GFP [35], pPIN1:PIN1:GFP [38], pPIN4:PIN4:GFP [39], pDR5:NLS:eYFP [38], pSMB:SMB:GFP [40] and pWOX5:ER:GFP [29] were introgressed by crossing into *blj jkd-4 scr-4* [15], *cbf3-1* (SAIL_244_D02 from NASC) and OX CBF3 A30 (CS69502 from NASC). OX CBF A30 was crossed to pPLT2:PLT2:eYFP and *shr-2* pSHR:SHR:GFP and examined in F1 generation. UAS::CBF3-YFP J0571 was introgressed by crossing into *blj jkd-4 scr-4* J0571 line [15]. Transgenic lines *shr-2 plt1-4 plt2-2* [41], *plt1-4 plt2-2* [9] and *shr-2* [42] were also used in this work.

### Genotyping

DNA extraction from F2 seedlings was used for genotyping by PCR using the following oligonucleotides: *blj* F 5′-CTTGAATCTCAAGCAGAAGCG-3′and R 5′-TCTTGGTTTCTTCTCTGCATCTC-3′; *jkd-4* F 5′-GGATGAAAGCAATGCAAAACA-3′ and R 5′-AATGTCGGGATGATGAACTCC-3′; s*cr-4* F 5′-TTATCCATTCCTCAACTTCAGT-3′and R 5′-TGGTGCATCGGTAGAAGAATT-3′ and *cbf3-1* F 5′-AGTCTTCTCTGGACACATGGC-3′ and R 5′-TCCATAACGATACGTCGTCATC-3′. T-DNA insertions were genotyped using the following pairs of primers: *blj* F 5′-CTTGAATCTCAAGCAGAAGCG-3′and R 5′-ACCCGACCGGATCGTATCGGT-3′; *jkd-4* F 5′-GGATGAAAGCAATGCAAAACA-3′ and R 5′-TCAAACAGGATTTTCGCCTGCT-3′; *cbf3-1* F 5′-AGTCTTCTCTGGACACATGGC-3′ and R 5′-TAGCATCTGAATTTCATAACCAATCTCGATACA C-3′. Mutation in s*cr-4* was genotyped by using the primers F 5′-CTTATCCATTCCTCAACTCTATT-3′and R 5′-TGGTGCATCGGTAGAAGAATT-3′ while absence of s*cr-4* mutation was detected using F 5′-TTATCCATTCCTCAACTTCAGT-3′and R 5′-TGGTGCATCGGTAGAAGAATT-3′.

### Microscopy analysis

Roots were stained with propidium iodide (10mg/mL. Sigma-Aldrich) and imaged using a TCS SP8 confocal microscope (Leica). Mature embryos were stained using the aniline blue staining method and scanned by confocal microscopy as described by [43]. Laser ablation of specific cells was performed with one to three pulses of 30 seconds with a UV laser (405nm) using FRAP mode of TCS SP8. Root meristem size measurements were performed as described by [21] using Image J program (Bethesda, MD, USA). Division of quiescent center (QC) cells upon laser ablation recovery was determined through confocal microscopy imaging of 1.5µm sections of the whole stem cell niche every 24 hours and scored as the number of QC cells generated by division of QC cells marked with *pWOX:ER:GFP* in the same root. pWOX5 marked cells was scored as the number of *pWOX5:ER:GFP* marked cells outside of QC using the same imaging approach. Quantification of fluorescent signal was performed using the hybrid detector counting mode and pixel density quantification in Leica LAS AF Lite software.

### Real time quantitative polymerase chain reaction (RT-qPCR) analysis

Root meristems were dissected at 6 day after imbibition using a micro scalpel (MICRO FEATHER) under a stereo microscope (Leica MZ95). RNA was isolated with the RNeasy isolation kit (Quiagen) and RNA integrity was measured with pico chip Bioanalyzer 2100. cDNA synthesis was performed with NYZ First-Strand cDNA Synthesis Kit (NYZ tech). RT-qPCR was performed in ECO Real-Time PCR System (Illumina) with MasterMix qPCR ROx PyroTaq EvaGreen dye (CMB) with ROX inner normalization. The following pairs of primers were used: AT3G07730 F 5′-TTGTTGTTGGATGGCAATCTG-3′and R 5′-AGTCTTGAGACATTGCATCAGT-3′ as control and *CBF3* F 5′-CAGTTTCAGTATAAGTGTGGGC-3′and R 5′-GCTGAATCGGTTGTTTCGGT-3′.

### Data collection and Statistical Analysis

For laser ablation experiments, at least 20 roots were assayed for each genetic background in two independent rounds. For root growth, meristem size and meristem differentiation measurements 60-90 seedlings of each genetic background in 2-3 independent rounds were used. SPSS Statistics 21 software (IBM) was used for statistical analysis. Normality and variance distributions were assayed using Kolmogorov-Smirnov and Levene tests, respectively. Significance in distributions of means was performed by one-way ANOVA analysis when samples to compare were <3 as indicated (case of one sample and control), or by General Linear Model (GLM) to assess for differences between genotypes, or when genotypes were compared at specific times, GLM was performed with a least significant difference (LSD) posthoc test.

### Microarray analysis

Microarrays correspond to reference GSE60157. Normalization and differentially expressed genes were calculated as previously described [15, 42], comparing set of microarrays only processed with the same Affymetrix kit and using the follwing cut-off settings: fold-change >±1.5, q-value < 10E-3 and expression value >1.

### Identification of transcription factors with enriched expression in the ground tissue

For identification of transcription factors with enriched expression in the ground tissue, Arabidopsis transcription factors in database PlantTFDB 3.0 [44] were required to be expressed in the ground tissue (J0571 cell type>1) and not be expressed (<1) in root-hair cells (COBL9 cell type), hairless cells (GL2 cell types), lateral root cap (J3411 cell type) nor columella (PET111 cell type) of the RootMap [45]. In addition, they were required to: a) not be expressed (<1) in the quiescent center (AGL42 cell type) nor the stele (WOL cell type) or b) be at least 2-fold-enriched in the ground tissue (J0571) in comparison with the quiescent center (AGL2) and the mean value of the cell types which make up the stele (J2661, JO121-pericycle, APL, S32-phloem, SUC2-companion cells, J2501, S4, S18-xylem). Intersection of all criteria to identify candidate genes was performed in Microsoft Access.

## Supporting information

Sanchez-Corrionero_SuppInformation

## ACKNOWLEDGMENTS

This work was supported by grants from Ministerio de Economía y Competitividad (MINECO) of Spain and ERDF BFU2013-41160-P and BFU2016-80315-P to M.A.M.-R. A.S.-C. was supported by a FPI contract from MINECO. P.P.-G. was supported by a Juan de la Cierva contract from MINECO and Programa Atraccion Talento from Comunidad Madrid. J.C was supported by a Juan de la Cierva contract from MINECO. We thank M. P. Gonzalez-Garcia, Ph.D, for reading and commenting this manuscript.

## SUPPORTING INFORMATION

**Figure S1. BLUEJAY, JACKDAW and SCARECROW maintain root growth capacity and meristem function. A**, root growth per day of WT, *scr4*, *blj jkd4*, *blj jkd4 scr4*, *plt1 plt2* and *shr plt1 plt2 seedlings*. mm: millimeters. Crosses and asterisks: statistically significant (p-value < 0.05 and < 0.001, respectively) by General Linear Model (GLM) and LSD post-hoc test as compared to the WT. a and b: statistically different (p-value < 0.05) when genotypes were compared separately for each day by GLM followed by LSD. Only differences between *blj jkd4 scr4* and *plt1 plt2* are indicated. **B**, confocal images of WT and *blj jkd4 scr4* mature embryos stained with aniline blue. e: endodermis, c: cortex, m: mutant layer, QC: quiescent center. Scale bar: 10 µm. **C**, meristem length of WT, *scr4*, *blj jkd4*, *jkd4 scr4*, *blj jkd4 scr4*, *plt1 plt2* and *shr plt1 plt2* at 3, 6, 9 and 13 dpi. Asterisk: statistically significant (p-value < 0.001) by GLM and LSD post-hoc test. **E**, confocal images of the RAM of sc*r4*, *blj jkd4*, *blj jkd4 scr4*, *plethora* (*plt)1 plt2* and *short-root-2* (*shr2) plt1 plt2*. White arrowheads indicate the end of the RAM. scale bars: 25 µm.

**Figure S2. BLUEJAY, JACKDAW *and* SCARECROW regulate proliferation and differentiation programs**. **A**, box plot representation of microarray expression values for cell cycle genes in the root apical meristem of WT and *blj jkd4 scr4*. **B**, confocal images showing *pCYCB1;1::CYCB1;1::GreenFluorescentProtein (GFP)* maximum projection of expression in the 3D root volume in WT and *blj jkd4 scr4* meristems at 7 dpi. Cell walls, in red (stained with propidium iodide -PI) are shown at the medium plane of the meristem. **C-D**, box-plot representations of mean-normalized expression values of **C**, repressed and **D**, activated genes in microarray experiments of *blj jkd4 scr4* meristems as compared to the WT.

**Figure S3. *SHORT-ROOT* and *PIN-FORMED1*, *2* and *4* auxin transporter genes are repressed in meristems of *blj jkd4 scr4 triple mutant.* A-B**, graphs showing microarray mean expression values for *SHR*, *PIN-FORMED 1* (*PIN1*), *PIN2* and *PIN2* genes in the RAM of WT and *blj jkd4 scr4*. p-values as indicated by mixed-model ANOVA analysis of replicates.

**Figure S4. DUO1-ACTIVATED ZINC FINGER 3 (DAZ3) and NUCLEAR FACTOR SUBUNIT B5 (NF-YB5) proteins are putatively mobile factors**. **A-B**, confocal images showing localization of YFP-tagged DUO1-ACTIVATED ZINC FINGER 3 (DAZ3) and NUCLEAR FACTOR SUBUNIT B5 (NF-YB5) in the *UAS/GAL4* system and enhancer trap line J0571 at 6 dpi. Expression of transgenes under *UAS* promoter in the J0571 line primarily corresponds to the ground tissue as indicated by localization of the GFP, although expression in the QC can be observed occasionally in several cells. Blue arrows: extra divisions in the endodermis. **C-D**, confocal images showing localization of YFP-tagged bHLH114 and RAVEN/INDETERMINATE DOMAIN 5 (RVN/IDD5) in the *UAS/GAL4* system and enhancer trap line J0571 at 6 dpi. **E**, confocal images of the meristem of *blj jkd4 scr4* J0571, *blj jkd4 scr4* J0571 *UAS::DAZ3::YFP* and *blj jkd4 scr4* J0571 *UAS::NF-YB5::YFP* at 12 dpi. **A-E**, Scale bar: 25µm.

**Figure S5. BLUEJAY, JACKDAW, SCARECROW regulate root regeneration**. **A**, graph showing incremental number of QC cells in WT and *blj jkd4 scr4* at 24h after laser ablation of stem cells above the QC. **B**, graph showing number of WOX5-marked cells in the 3-D root meristem volume at 24 and 48h after stem cell ablation in WT and *blj jkd4 scr4*.

