## Supplementary material for "ROOT PATTERNING AND REGENERATION ARE MEDIATED BY THE QUIESCENT CENTER AND INVOLVE BLUEJAY, JACKDAW AND SCARECROW REGULATION OF VASCULATURE FACTORS": Sanchez-Corrionero_SuppInformation

**Supplementary Table 1. Transcription factors in the SHR network with enriched expression in the ground tissue. Related to Figure 4**

| ***Locus*** | ***Name*** | ***J0571 / UAS construct*** | ***Mobile*** |
|---|---|---|---|
| AT1G14580 | BLJ / IDD6 | YES | NO |
| AT2G02070 | RVN / IDD5 | YES | NO |
| AT2G46680 | ATHB7 | NO | - |
| AT2G47810 | NF-YB5 | YES | YES |
| AT4G05170 | BHLH114 (E29) | YES | NO |
| AT4G21340 | BHLH103 / B70 (E30) | YES | NO |
| AT4G25480 | CBF3 / DREB1A | YES | YES |
| AT4G35700 | DAZ3 | YES | YES |
| AT5G58620 | TZF9 | NO | - |

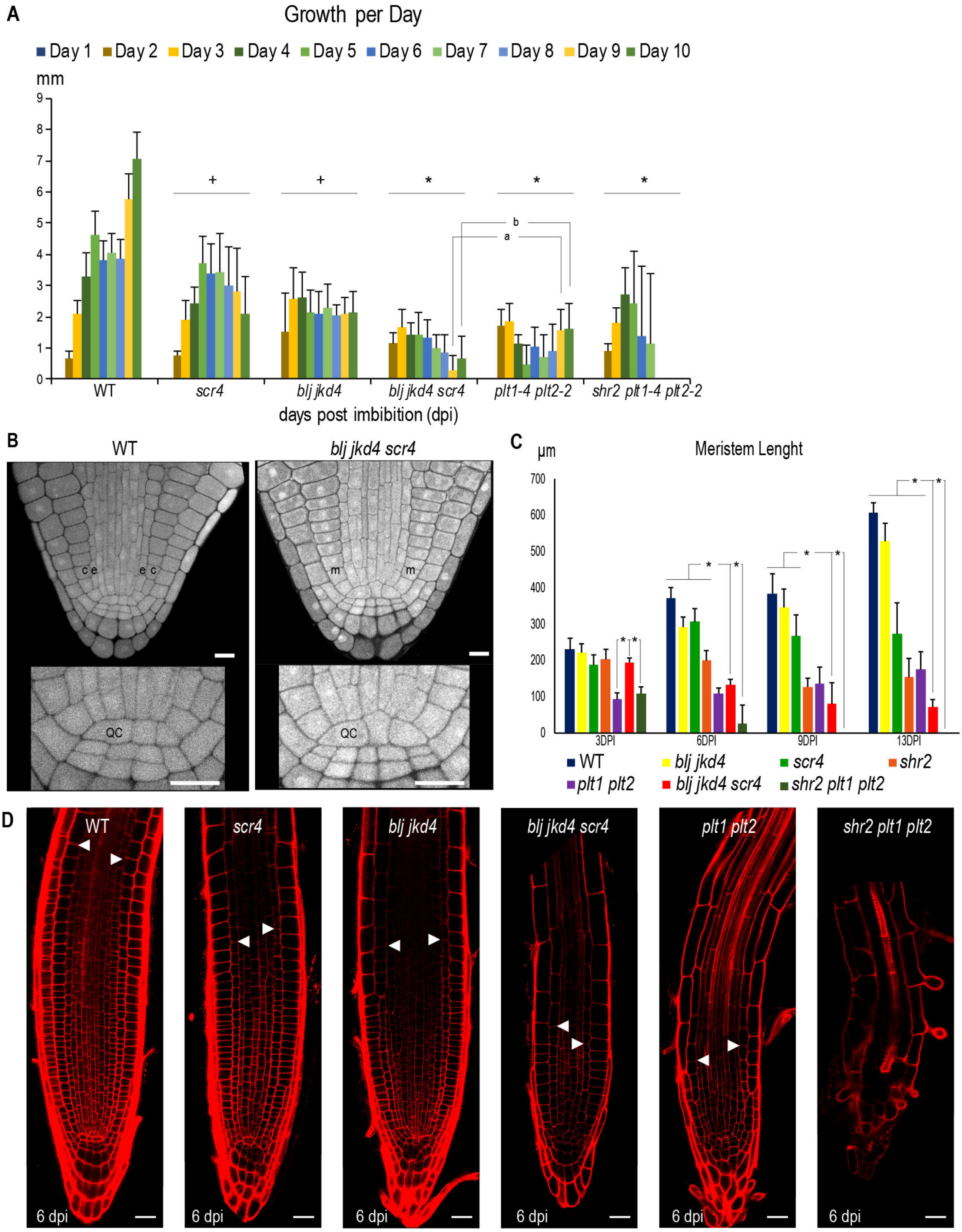


**Figure S1. BLUEJAY, JACKDAW and SCARECROW maintain root growth capacity and meristem function. A**, root growth per day of WT, *scr4*, *blj jkd4*, *blj jkd4 scr4*, *plt1 plt2* and *shr plt1 plt2 seedlings*. mm: millimeters. Crosses and asterisks: statistically significant (p-value < 0.05 and < 0.001, respectively) by General Linear Model (GLM) and LSD post-hoc test as compared to the WT. a and b: statistically different (p-value < 0.05) when genotypes were compared separately for each day by GLM followed by LSD. Only differences between *blj jkd4 scr4* and *plt1 plt2* are indicated. **B**, confocal images of WT and *blj jkd4 scr4* mature embryos stained with aniline blue. e: endodermis, c: cortex, m: mutant layer, QC: quiescent center. Scale bar: 10 µm. **C**, meristem length of WT, *scr4*, *blj jkd4*, *jkd4 scr4*, *blj jkd4 scr4*, *plt1 plt2* and *shr plt1 plt2* at 3, 6, 9 and 13 dpi. Asterisk: statistically significant (p-value < 0.001) by GLM and LSD post-hoc test. **E,** confocal images of the RAM of s*cr4*, *blj jkd4*, *blj jkd4 scr4*, *plethora* (*plt*)*1 plt2* and *short-root-2* (*shr2) plt1 plt2*. White arrowheads indicate the end of the RAM. scale bars: 25 µm.

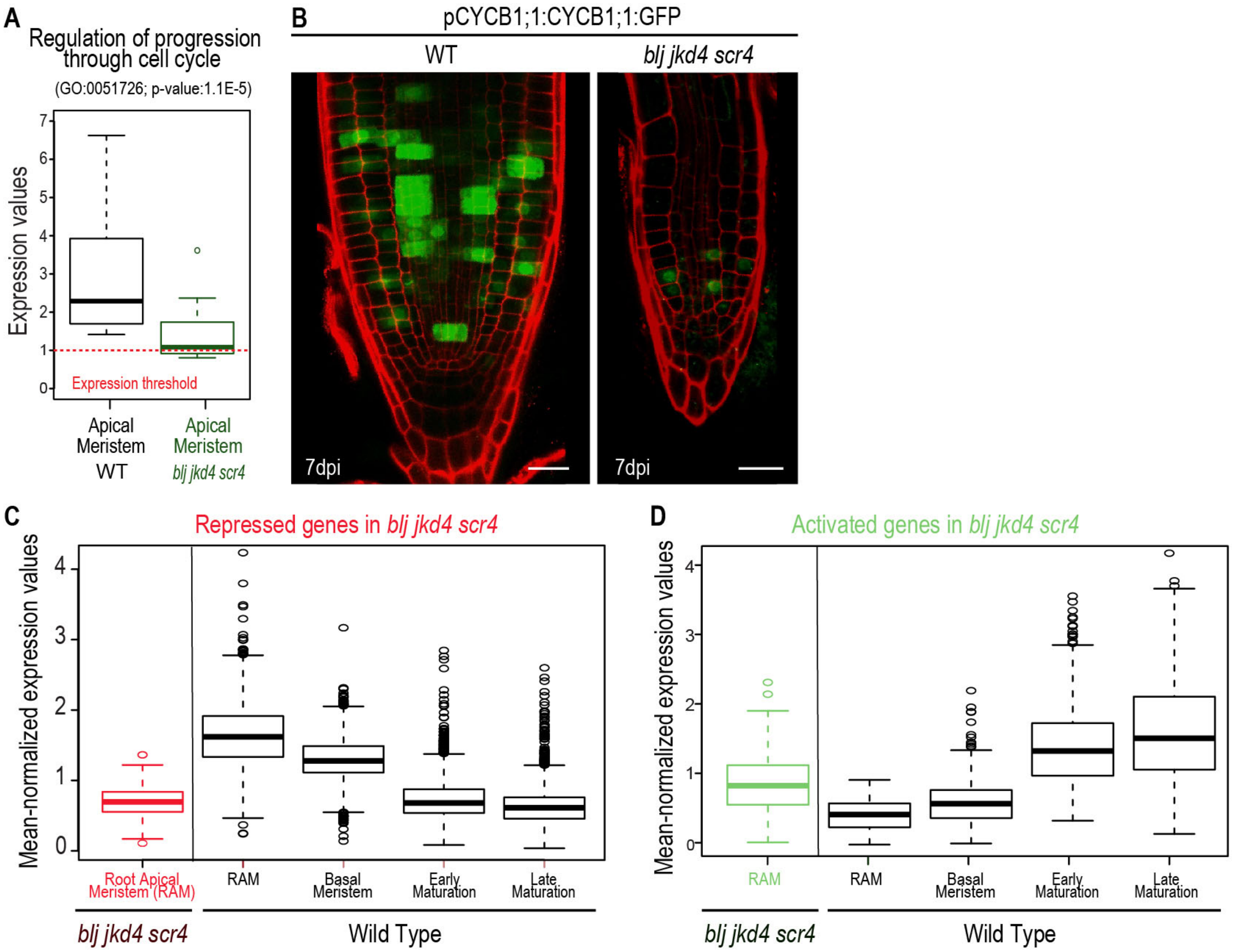


**Figure S2. BLUEJAY*, *JACKDAW *and* SCARECROW regulate proliferation and differentiation programs**. **A**, box plot representation of microarray expression values for cell cycle genes in the root apical meristem of WT and *blj jkd4 scr4*. **B,** confocal images showing *pCYCB1;1::CYCB1;1::GreenFluorescentProtein (GFP)* maximum projection of expression in the 3D root volume in WT and *blj jkd4 scr4* meristems at 7 dpi. Cell walls, in red (stained with propidium iodide -PI) are shown at the medium plane of the meristem. **C-D**, box-plot representations of mean-normalized expression values of **C,** repressed and **D,** activated genes in microarray experiments of *blj jkd4 scr4* meristems as compared to the WT.

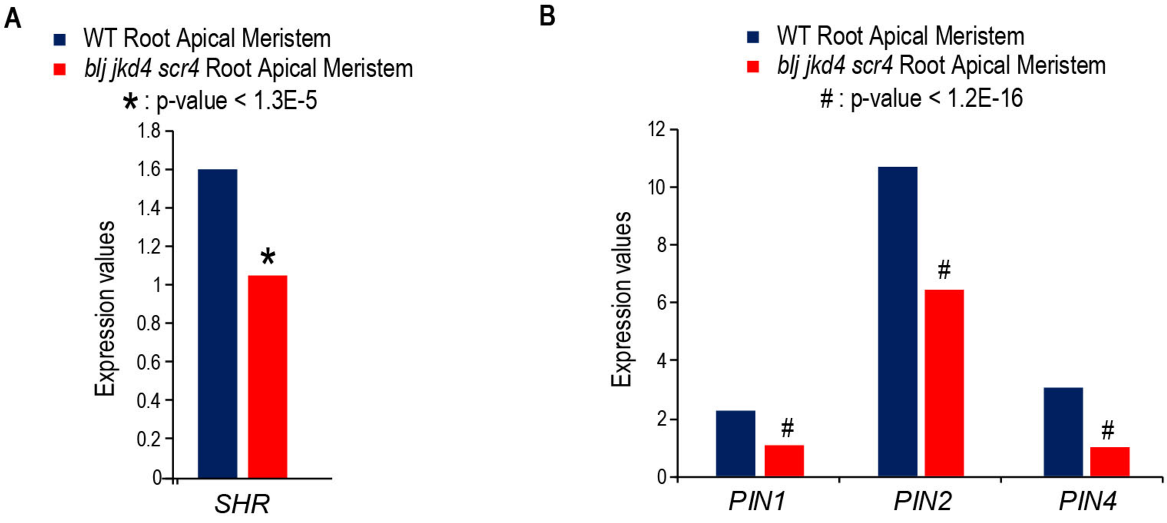


**Figure S3. *SHORT-ROOT* and *PIN-FORMED1*, *2* and *4* auxin transporter genes are repressed in meristems of *blj jkd4 scr4 triple mutant.*** **A-B**, graphs showing microarray mean expression values for *SHR*, *PIN-FORMED 1* (*PIN1*), *PIN2* and *PIN2* genes in the RAM of WT and *blj jkd4 scr4*. p-values as indicated by mixed-model ANOVA analysis of replicates.

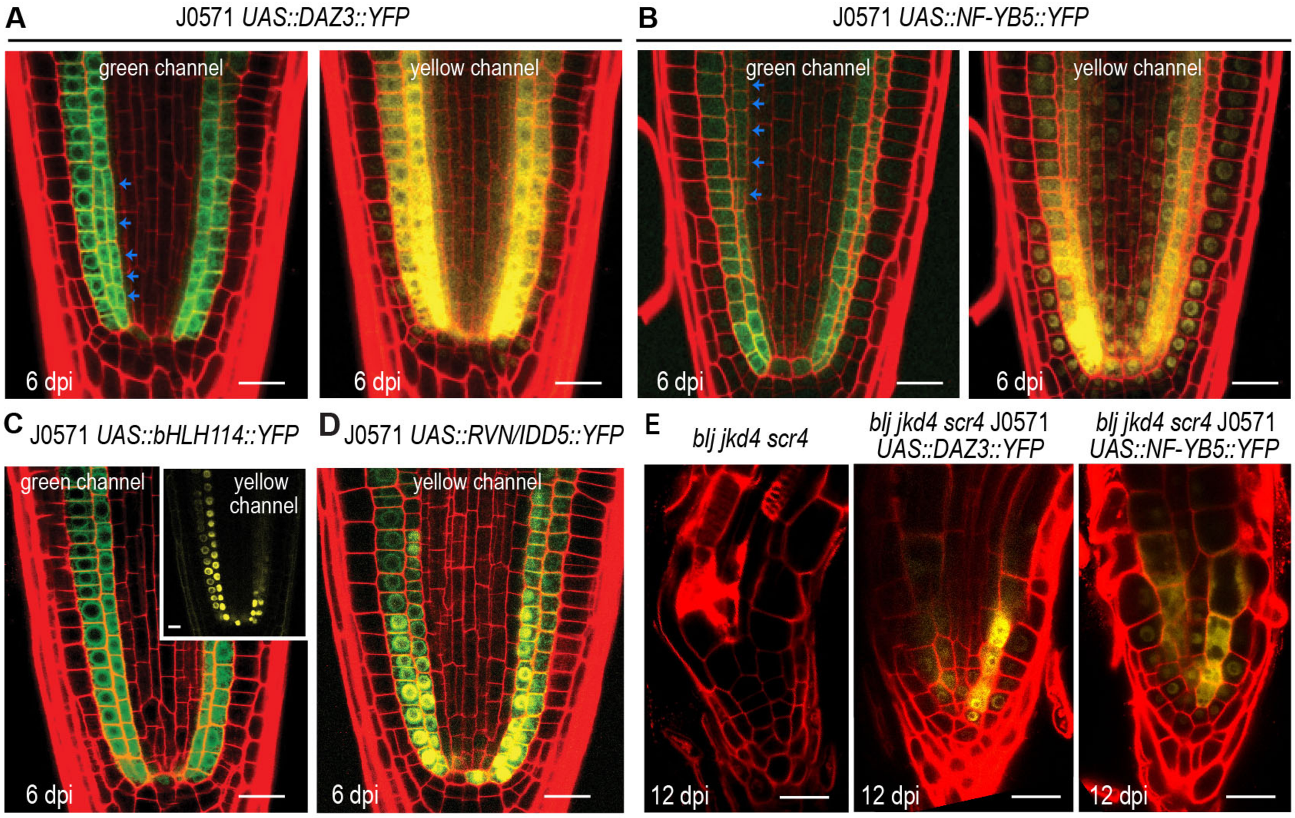


**Figure S4. DUO1-ACTIVATED ZINC FINGER 3 (DAZ3) and NUCLEAR FACTOR SUBUNIT B5 (NF-YB5) proteins are putatively mobile factors**. **A-B**, confocal images showing localization of YFP-tagged DUO1-ACTIVATED ZINC FINGER 3 (DAZ3) and NUCLEAR FACTOR SUBUNIT B5 (NF-YB5) in the *UAS/GAL4* system and enhancer trap line J0571 at 6 dpi. Expression of transgenes under *UAS* promoter in the J0571 line primarily corresponds to the ground tissue as indicated by localization of the GFP, although expression in the QC can be observed occasionally in several cells. Blue arrows: extra divisions in the endodermis. **C-D**, confocal images showing localization of YFP-tagged bHLH114 and RAVEN/INDETERMINATE DOMAIN 5 (RVN/IDD5) in the *UAS/GAL4* system and enhancer trap line J0571 at 6 dpi. **E**, confocal images of the meristem of *blj jkd4 scr4* J0571, *blj jkd4 scr4* J0571 *UAS::DAZ3::YFP* and *blj jkd4 scr4* J0571 *UAS::NF-YB5::YFP* at 12 dpi. **A-E**, Scale bar: 25µm.

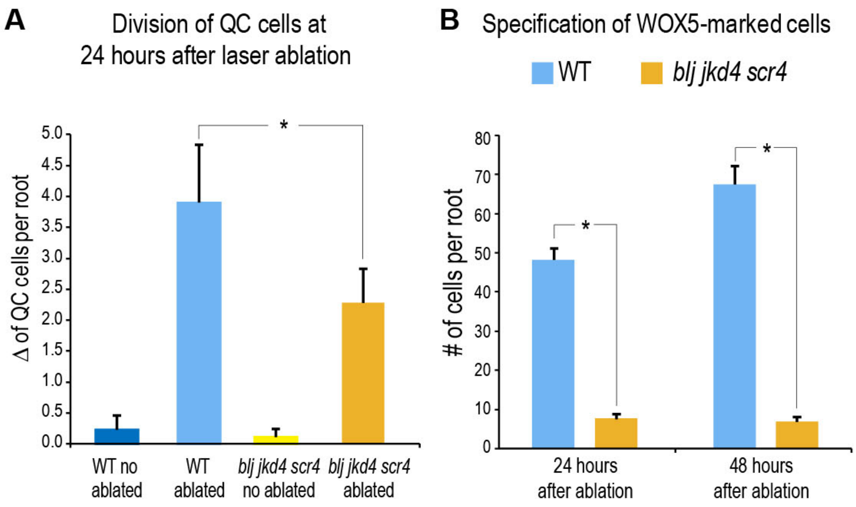


**Figure S5. BLUEJAY, JACKDAW, SCARECROW regulate root regeneration**. **A**, graph showing incremental number of QC cells in WT and *blj jkd4 scr4* at 24h after laser ablation of stem cells above the QC. **B**, graph showing number of WOX5-marked cells in the 3-D root meristem volume at 24 and 48h after stem cell ablation in WT and *blj jkd4 scr4*.
